# Insights into Drug Cardiotoxicity from Biological and Chemical Data: The First Public Classifiers for FDA DICTrank

**DOI:** 10.1101/2023.10.15.562398

**Authors:** Srijit Seal, Ola Spjuth, Layla Hosseini-Gerami, Miguel García-Ortegón, Shantanu Singh, Andreas Bender, Anne E. Carpenter

## Abstract

Drug-induced cardiotoxicity (DICT) is a major concern in drug development, accounting for 10-14% of postmarket withdrawals. In this study, we explored the capabilities of various chemical and biological data to predict cardiotoxicity, using the recently released Drug-Induced Cardiotoxicity Rank (DICTrank) dataset from the United States FDA. We analyzed a diverse set of data sources, including physicochemical properties, annotated mechanisms of action (MOA), Cell Painting, Gene Expression, and more, to identify indications of cardiotoxicity. We found that such data, including protein targets, especially those related to ion channels (such as hERG), physicochemical properties (such as electrotopological state) as well as peak concentration in plasma offer strong predictive ability as well as valuable insights into DICT. We also found compounds annotated with particular mechanisms of action, such as cyclooxygenase inhibition, could distinguish between most-concern and no-concern DICT compounds. Cell Painting features related to ER stress discern the most-concern cardiotoxic compounds from non-toxic compounds. While models based on physicochemical properties currently provide substantial predictive accuracy (AUCPR = 0.93), this study also underscores the potential benefits of incorporating more comprehensive biological data in future DICT predictive models. With the availability of - omics data in the future, using biological data promises enhanced predictability and delivers deeper mechanistic insights, paving the way for safer therapeutic drug development. All models and data used in this study are publicly released at https://broad.io/DICTrank_Predictor

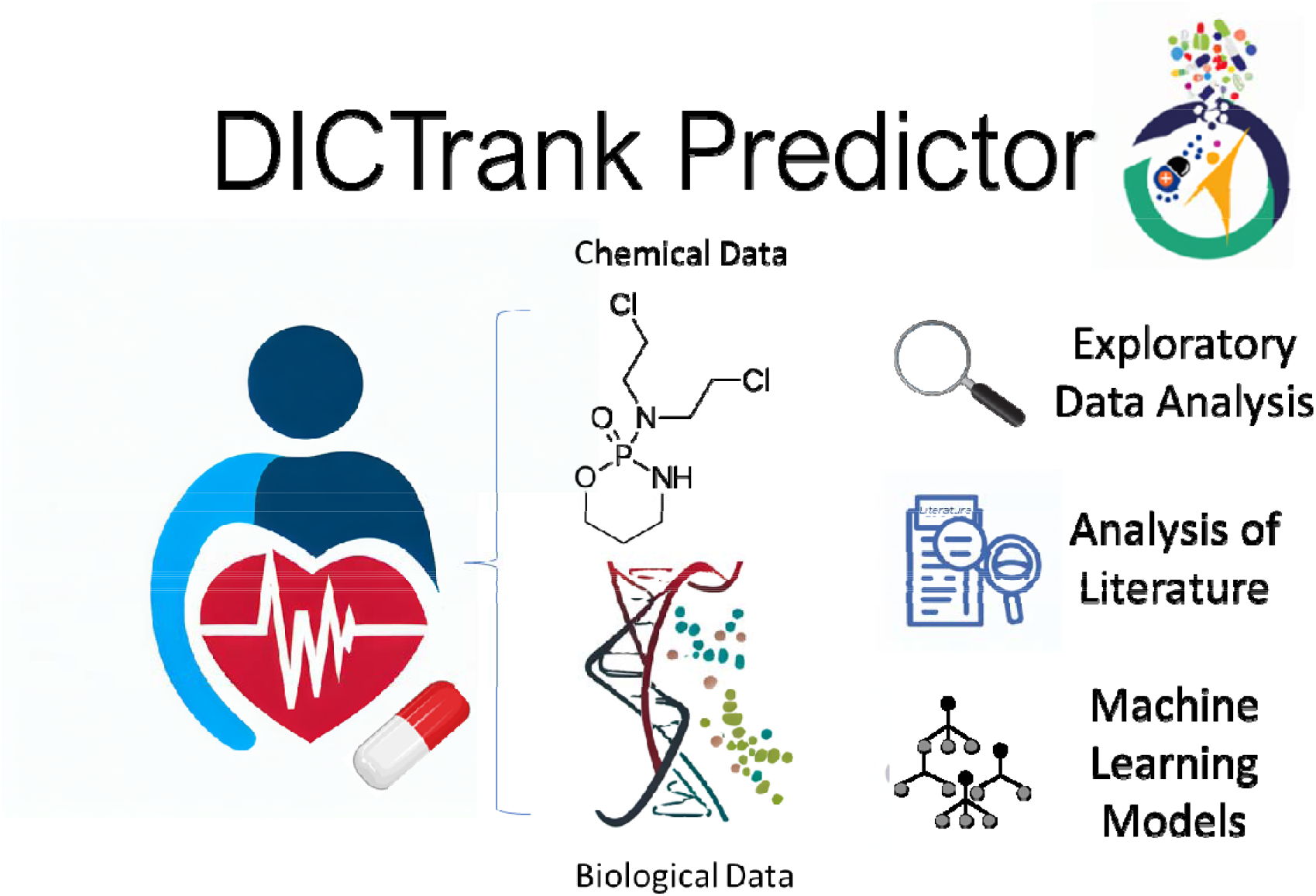

## Introduction

Drug-induced cardiotoxicity (DICT) is a leading cause of drug withdrawals during post-market surveillance. One study showed that 10% of withdrawals in the last 4 decades were due to cardiovascular safety concerns, including previously successful therapeutics such as rofecoxib, tegaserod, sibutramine, and rosiglitazone.^1^ Another study found that cardiotoxicity was the third most common reason for adverse drug reactions and accounted for 14% of withdrawals.^2^ Worryingly, the rate of DICT-related withdrawals may even be increasing, accounting for 17 out of 38 cases among drugs approved between 1994 and 2006.^1, 3^

DICT is associated with both functional damage such as arrhythmia, which alters mechanical function, and structural damage such as morphological damage in cardiomyocytes; functional damage and structural damage in the heart can be interrelated, where one may precipitate the other.^4^ DICT can be attributed to several underlying mechanisms affecting myocardial functions and viabilities.^5^ Some drugs, such as anthracyclines, inflict direct myocyte injury via reactive oxygen species production and compromising DNA replication.^6^ Electrophysiological disruptions, for example, measured in the hERG potassium channel blockers, can lead to arrhythmias by causing QT interval prolongation.^7^ Cardiac energy demands can be affected by drugs that interfere with mitochondrial functionality.^1^ Drugs may also adversely influence vascular supply, inducing ischemic conditions.^8^ Intracellular calcium regulation for cardiomyocyte activity can also disrupt its homeostasis, resulting in contractile and rhythm abnormalities.^9^ Furthermore, alterations in growth factors and cytokine balances can induce cardiac conditions like fibrosis, and immunologic drug reactions can also cause cardiotoxicity.^10, 11^ Several neurohormonal pathways also offer indirect routes for drug-induced cardiac stress.^12^ Notably, a single drug might induce cardiotoxicity via multiple mechanisms, and individual patients’ responses (which can often manifest as side effects) can be modulated by genetics, concurrent health conditions, and other medications.^13^

To move beyond a limited focus on specific adverse reactions or related proxy assays for cardiotoxicity, the FDA recently released the Drug-Induced Cardiotoxicity Rank (DICTrank) that categorizes drugs based on their risk of causing cardiotoxicity.^14^ Similar to the DILIrank data for liver injury^15^, the DICTrank system uses FDA drug labeling to comprehensively categorize 1,318 human drugs into four DICT Concern categories based on their potential risk for cardiotoxicity: (1) Most-DICT-Concern, (2) Less-DICT-Concern, (3) No-DICT-Concern and (4) Ambiguous-DICT-Concern. The DICTrank dataset was generated with an expertise review from the FDA, keyword searches, and manual curation of FDA labeling documents as well as data from clinical trials, post-marketing, and literature surveys.

Predictive models for drug-induced cardiotoxicity (DICT) could save considerable time, resources, and human suffering, with the ultimate goal of preventing adverse events in clinical trials and the post-market stage. However, predicting any *in vivo* effect is not a trivial classification task, and most predictive models are built on proxy endpoints (which are often reduced to binary endpoints) without taking into account *in vivo* parameters such as pharmacokinetic parameters.^16^ While no models for DICTrank have been publicly available yet to the best of our knowledge, various studies have predicted proxy *in vitro* assays or side effect data from SIDER (Side Effects Resource), some of which are related to cardiotoxicity.^17^ Studies focusing on side effects and proxy targets (such as hERG) are reasonable given that compounds that have cardiac-related indications are more likely to show related side effects as well or activity on ion channels.^7^

Previously it was shown that adverse events data and biological data can be used for identifying mechanism hypotheses leading to cardiotoxicity.^18^ Wang et al used LINCS L1000 gene expression features to predict a wide range of drug-induced adverse events from the SIDER dataset.^19^ Particularly for acute myocardial infarction, models developed achieved an AUC-ROC of 0.84 when using chemical structural data and 0.76 when using Gene Ontology annotations (compared to 0.5 for random models). Gelano et al used a matrix decomposition algorithm to predict side effect frequencies for drugs and provide biologically interpretable insights.^20^ MoleculeNet predictions for SIDER side effects, trained on chemical structure data, range from 0.65 to 0.70 AUC-ROC when using a bypass network, a modified version of a multi-task network.^21^

Most predictive models above were built on chemical structure data as input features. Although certain structural motifs or patterns in a molecule can be indicative of toxic properties and analyzing the chemical structure can flag potential cardiotoxic compounds, such models are often limited in their applicability domain, that is, their accuracy is limited to the chemical space of the training data, and they fail to generalize to markedly different chemical structures. Novel chemical and biological data have been previously used to evaluate side effects in general from the SIDER dataset.^22^ Previous studies have shown that Random Forest models trained on a combination of biological, chemical, and phenotypic features achieved an AUCPR of 0.76 for cardiac disorders.^23^

With the availability of the new DICTrank dataset, we used a novel multi-faceted approach using both chemical and biological data (that considers a multitude of possible mechanisms that can lead to DICT) intending to better understand and make mechanistic insights into a drug’s cardiac safety profile. We evaluated a wide range of chemical and biological information, as shown in Figure 1, to determine which feature space is most predictive of DICTrank and evaluated these feature spaces to build the first predictive models of DICTrank using machine learning. Biological data sources included Cell Painting, gene expression, and Gene Ontology,^24–28^ as well as bioactivity, and annotated mechanisms of action (MOA)^29^ and pharmacokinetic parameters for the peak unbound and total concentration of a drug molecule in plasma^30^; these offer an alternate feature space to chemical space.^31^ We aimed to glean insights from which chemical and biological data best capture the carefully curated manual annotations in the DICTrank data. Incorporating data from all these sources as feature spaces for predictive models allows for a multifaceted assessment of a drug’s potential cardiotoxicity, potentially enhancing the model’s accuracy and reliability. Overall, the use of biological data sources along with chemical data improved detection and offered mechanistic insights into the cardiotoxicity of compounds. The models based on chemical structures and physicochemical characteristics are readily accessible for direct use on <> (owing to the constrained availability of public data for other feature types). All code and data for all models can be found on GitHub (https://github.com/srijitseal/DICTrank) for local implementation with further details on https://broad.io/DICTrank_Predictor.

**Figure 1:**
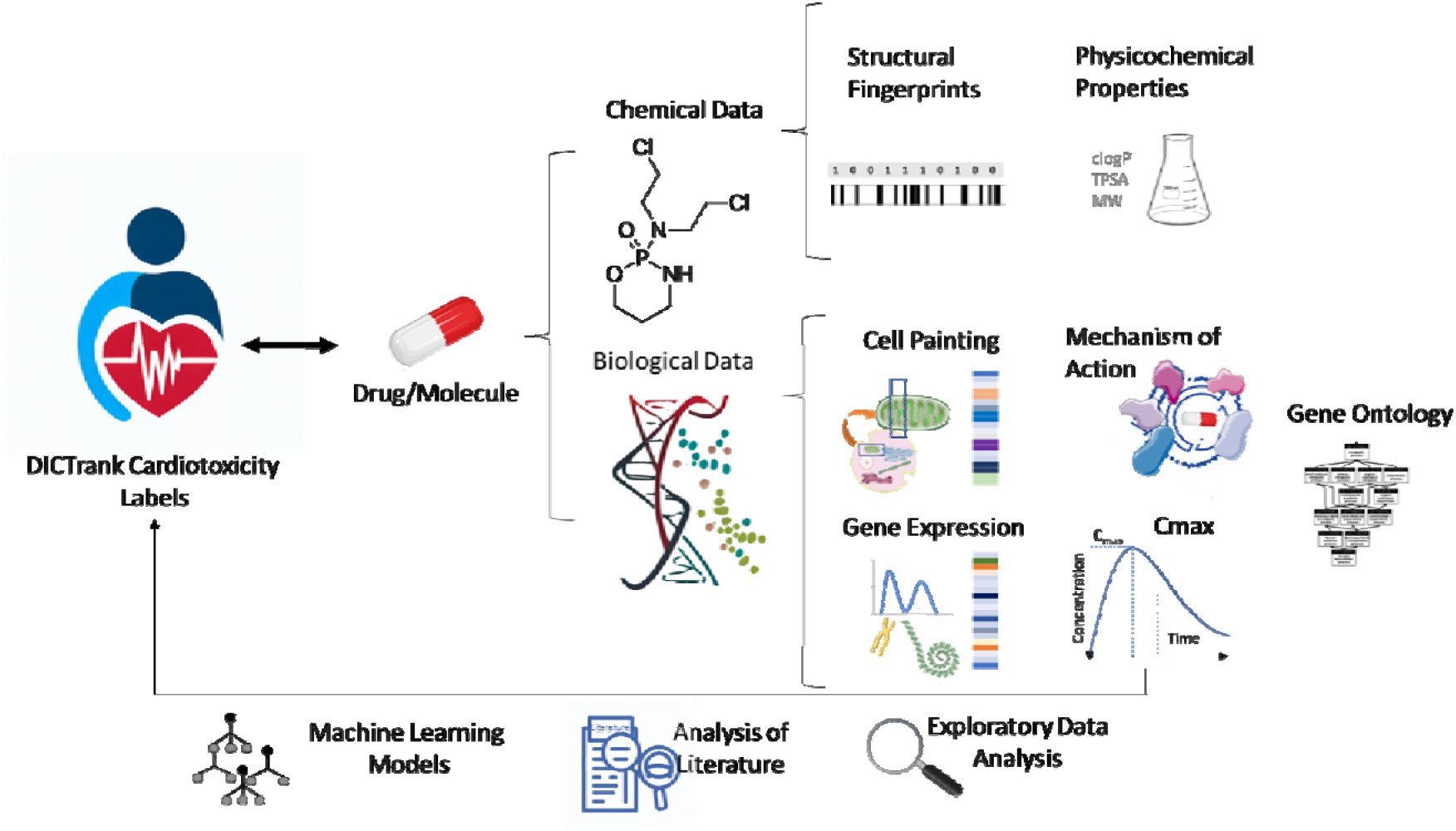
Chemical and biological data sources were used in this study to perform exploratory data analysis on DICTrank and training predictive models (further details in Table 2).

## Methods

### Data sources

We obtained the DICTrank dataset, as released by Qu et al. which includes comprehensive DICTConcern categories for a diverse set of over 1300 drugs.^14^ The SIDER database, a pharmacovigilance resource, contained associations for drugs with side effects.^21, 32^ We used data from cardiac disorders from the SIDER dataset to compare concordance with DICTrank and enrich the dataset as described later. To gain insights into the mechanisms of action (MOA) of various drugs, we assessed relevant data from the Drug Repurposing Hub^33^ which contained information on 6777 drugs for 1130 MOAs and 2183 known targets. To explore the potential targets of drugs, we incorporated the CellScape target predictions on inhibition/antagonism for 2,094 targets at four concentrations (0.1, 1, 10, and 100 uM).^34^ We used morphological profiles from the Cell Painting assay^24^ which considers the impact of drugs on cellular morphology and function. This dataset contained a range of circa 1700 morphological features for over 15,000 compound perturbations. We obtained gene expression data from LINCS L1000 data which contains over 19,000 drugs as described in Wang et al.^19^ This study utilized gene expression features derived from LINCS L1000^25^ transcriptomic data, capturing changes in 978 landmark genes across diverse human cell lines in response to compound perturbations. Gene Ontology-transformed expression features^26^, which encode biological processes involved with gene expressions affected by the compound perturbations, were extracted from a dataset containing 4,438 annotated features linked to these compounds in the study.^19^ The analysis by Wang et al prioritized the strongest signatures across cell line, concentration, and time point for each compound using Characteristic Direction (CD) and evaluated the enrichment across various gene set libraries via Principal Angle Enrichment Analysis (PAEA).^35^ Finally we used pharmacokinetic data, specifically the maximum unbound and total concentrations (Cmax) of 758 drugs in the bloodstream, as compiled by Smith et al.^30^ This dataset contains Cmax (unbound) for 534 compounds and Cmax (total) for 749 compounds.

### Standardization of the SMILES

For each dataset, we standardized chemical SMILES iteratively using RDKit^36^ and MolVS^37^ functionalities. This includes steps for InChI transformation, molecular cleanup, charge neutralization, tautomer normalization, and final standardization. We carried out up to five iterations of the standardization until a standardized SMILES was finalized, otherwise, we chose the most common SMILES from the counter. Finally, the molecule was protonated at pH 7.4 using DimorphiteDL to reflect its likely state at physiological pH.^38^ Hence, we obtained a standardized SMILES and a standardized InChI.

### Preprocessing data

For the DICTrank dataset, we binarized the dataset considering DICT no-concern as 0 and less- and most-concern as 1 as DICTrank labels for machine learning classifiers. We removed compounds that were ambiguous and treated a compound as toxic if there was at least one record of toxicity among duplicates. For the SIDER dataset, we removed duplicate standardized smiles, and similar to the above, labeled a compound as toxic if there was at least one evidence of toxicity among the duplicates. Labels from both SIDER and DICTrank are described in Table 1.

**Table 1:** Distribution of compound toxicity labels related to cardiotoxicity/ cardiac disorders for all unique compounds from each of the datasets used in this study.

| Dataset | Label | Number of Toxic Compounds | Number of Non-Toxic Compounds | Description |
| --- | --- | --- | --- | --- |
| SIDER | Cardiac disorders (binary) | 829 | 360 (Absence of Evidence) | Recorded adverse drug reactions from marketed medicines. |
| DICTrank | DICT Concern Category (Categorical) | Most: 299, Less: 443, | No: 278 (Evidence of Absence) | A ranking system from DICTrank that categorizes drugs according to risk for cardiotoxicity. |
|  | DICTrank label (binary) | 742 | 278 (Evidence of Absence) | Binarized labels obtained from DICT Concern categories used in this study |

**Table 2:** The description of various feature spaces used in this study.

| Feature Space | Dimensions after feature selection (where applicable) | Description | Signal Expected | Source |
| --- | --- | --- | --- | --- |
| Chemical Structure | 2048-bit vector | ECFP4 (Morgan) fingerprints representing chemical structures | Distinctive patterns of chemical bonding and arrangement | 52 |
| Physicochemical properties (Mordred Descriptors) | 1038 2-D descriptors | Properties such as lipophilicity, solubility, molecular weight, ionizing potential, etc. | Properties that are associated with negative impacts on ion channels in the heart | 41 |
| MOA dataset | 264 binary encoded MOAs + 551 known targets | Annotations for mechanism of action and known targets based on knowledge. | Mechanism of action for drugs that inhibit certain ion channels | 29 |
| CellScape Target Prediction dataset | 1893 predictions for 817 unique targets and concentration combinations (0.1, 1, 10, and 100uM) | Predicted protein target for inhibition/antagonism; does not consider the functionality; prediction is based on chemical structure; updated algorithm from PIDGINv4<sup>43</sup> | Understanding how a drug interacts with various biological targets (not just its primary target) can provide insights into potential off-target effects | 34,43 |
| Cell Painting | 1783 features | Morphological changes in U2OS cells by a chemical perturbation, using a 5-channel fluorescence microscopy assay | Morphological changes in cells that reflect basic biological processes | 24,53 |
| Gene Expression | 978 features | Transcriptomic changes in response to chemicals using the L1000 assay | Upregulation or downregulation of genes associated with cardiac stress, apoptosis in cardiac cells, or ion channel function | 19,25 |
| Gene Ontology | 4438 annotations | Gene Ontology manual annotations based on collective knowledge | Understanding the biological processes, cellular components, and molecular functions affected, e.g., related to cardiac function, cardiac muscle tissue development, or ion homeostasis | 19,26 |
| Cmax | 2 features | The maximum total and unbound concentration of a drug in plasma | High Cmax would indicate a high risk of cardiotoxicity | 30 |

For the Cell Painting, Gene Expression, and Gene Ontology datasets, we use median cell profiles over standardized SMILES obtaining two datasets: 1783 Cell Painting features for 15,406 compounds, and Gene Expression features for 978 landmark genes and 4,428 Gene Ontology annotations for 9132 compounds. For the MOA dataset, we used one hot encoding of given annotations for compounds, which effectively gives us data for evidence of the presence of MOA/known targets and the absence of evidence. We used a variance threshold of 0.001 to identify and remove low-variance features reducing the dimensionality to 264 MOA and 551 known target features with significant variability. All datasets are released publicly at figshare (10.6084/m9.figshare.24312274) and https://broad.io/DICTrank_Predictor.

### Analyzing chemical space overlap between SIDER and DICTrank

We used standardized InChI to calculate the overlap between SIDER and DICTrank datasets. We assessed the physicochemical space using a t-distributed stochastic neighbor embedding (TSNE; as implemented in scikit-learn^39^) for six physicochemical properties, namely, molecular weight, topological polar surface area, number of rotatable bonds, hydrogen bond donors and acceptor, and the computed logarithm of the partition coefficient. To analyze the chemical space we used a Principal Component analysis (PCA) of the FragFP fingerprints from DataWarrior^40^, which in our experience works better with a higher explained variance in the plot of the principal component analysis compared to Morgan fingerprints.

### Structural and physicochemical features

For structural features, we used 2048-bit Morgan Fingerprints as implemented in RDKIT.^36^ For chemical compounds, we computed 1579 descriptors using Mordred.^41^ These physicochemical descriptors are derived from 2D representations of compounds, that is, we did not consider 3D descriptors. We removed the descriptors that failed to compute and finally obtained 1038 2-D physicochemical descriptors, and these were used for the machine learning models. For the analysis of feature distributions, we used the full set of 208 RDKit descriptors, (which are better interpretable compared to Mordred descriptors) as defined in the Descriptors module.^36^

### Predicted targets from CellScape

To derive predicted molecular targets for compounds, we utilized the commercially available CellScape target prediction package (Ignota Labs, 2023).^34^ This package applies models trained on a mixture of publicly available and proprietary bioactivity data (primarily inhibitory/antagonistic mechanisms) at 0.1, 1, 10, and 100uM with chemical structural features to output a probability score (between 0 and 1) of predicted activity for 2,094 distinct human targets. Although not used in this study, publicly available target prediction alternatives are also available such as PIDGINv4^42, 43^ and swisstargetpred^44^. We provide the computed CellScape features for compounds in the DICTrank dataset publicly via figshare (10.6084/m9.figshare.24312274) and https://broad.io/DICTrank_Predictor.

### Substructure analysis and retrospective analysis of DrugBank

For substructure analysis, we used SARpy^45^ on the DICTrank dataset, in a method similar to the one applied by Hemmrich et al.^46^. SARpy uses a recursive algorithm for fragmentation. We used two distinct settings for analysis: (1) using both toxic and non-toxic compounds and (2) using only toxic compounds to yield the desired substructures. For both settings, we confined the fragment size within a range of two to 18 atoms, with a minimum occurrence of five times. Furthermore, the positive predictive value (PPV) was adjusted to minimize false negatives. We combined structural alerts from both settings and quantified the frequency of these fragments within the entirety of the DICTrank dataset. We eliminated fragments with a PPV below 0.5. We then manually assessed the remaining fragments, for example removing those having four or fewer atoms, removing substructures like benzene, etc. to obtain 58 structural alerts.

We analyzed all compounds in DrugBank^47^ for the presence of structural alerts from the above to evaluate the risk of the chemical space of drugs for cardiotoxicity. We only used the compounds that did not overlap with the DICTrank dataset for this retrospective analysis to avoid information leaks. We annotated these compounds with labels for cardiac disorders from SIDER and disease area labels from the MOA dataset. We then checked for the presence of structural alerts among the subset of compounds that are currently approved, investigational, experimental, and/or withdrawn drugs.

### Analysis of chemical and biological data for differences in feature distribution for DICTrank compounds

We detected features that are predictive of highly cardiotoxic compounds. In order to do this, we detected features for each chemical and biological dataset that had a significant difference in the distribution for the DICT concern categories. For categorical features (SIDER, MOA annotations, and some of the 208 RDKit descriptors), we employed the chi-squared test (as implemented in SciPy^48^) to evaluate the association between categorical variables. We used a contingency table delineating the frequency distribution for each combination of category values. The chi-squared test yielded a statistical value alongside a corresponding p-value. For continuous features (as in the Cell Painting, Gene Expression, Gene Ontology datasets, and some of the 208 RDKit descriptors), we chose the Kruskal-Wallis test (as implemented in SciPy^48^) for evaluating the DICT-Concern labels since it is suited for comparisons involving three or more independent groups. Conversely, when comparing two classes, pairwise, the Mann-Whitney U test (as implemented in SciPy^48^) was used which is adept at discerning differences in distributions between two independent samples. Both tests yield a statistic value alongside its corresponding p-value. For both total unbound/plasma concentrations, as in the Cmax dataset, we used the Mann-Whitney U test to compare the distribution of Cmax among each DICT concern class and the DICTrank label.

### Enriching DICTrank compounds with SIDER compounds

We next determined the overlap of compounds (and the concordance in their labels) in the DICTrank dataset with the compounds in SIDER labeled with “Cardiac disorders” using the standardized InChI yielding 776 compounds in common. We next enriched DICTrank with SIDER giving a preference to the DICTrank label in case of a conflict. In this manner, we obtained three datasets besides the DICTrank dataset with the distribution of toxic/non-toxic compounds given in Supplementary Table S1. These are (1) DICTrank, (2) DICTrank enriched with cardiotoxic compounds from SIDER, (3) DICTrank enriched with non-cardiotoxic compounds from SIDER and (4) DICTrank enriched with all compounds from SIDER.

### Training predictive models for DICTrank

We trained eleven Random Forest models, each using the following features (as listed in Table 2): (1) Structural fingerprints, (2) Mordred descriptors, (3) MOA labels, (4) MOA labels along with total Cmax, (5) MOA labels along with unbound Cmax, (6) CellScape predicted protein targets, (7) CellScape predicted protein targets along with total Cmax, (8) CellScape predicted protein targets along with unbound Cmax, (9) Cell Painting features, (10) Gene Expression features, and (11) Gene Ontology features.

The training data available for these models depended on the number of compounds for which data was available and varied as given in Supplementary Table S1. As the external test set, we aimed to keep that fixed for a fair evaluation depending on available data as shown in Supplementary Table S2. For models not using Cmax data (where overlaps were larger and hence more data was available), we randomly selected 90 compounds (8.8% of the dataset, 65 cardiotoxic and 21 non-toxic) for which all annotations of feature spaces were available (as described in Supplementary Table S1). These 90 compounds struck a similar balance of DICT concern categories (most: 39, less: 26, and no: 25) as the original DICTrank dataset. For models using total Cmax data, we used the same external test set comprising 90 compounds since total Cmax data was available for these compounds. However, for models using unbound Cmax data (which had smaller overlaps compared to the above), we used a subset of 78 compounds (57 cardiotoxic and 21 non-toxic) as the external test set as shown in Supplementary Table S2.

Among the models that relied on -omics data (Cell Painting, Gene Expression, and Gene Ontology) we checked for each training compound whether a profile (feature set) was available. If there was no profile available in the respective datasets, we calculated the median profile of all compounds in the original dataset using a *v*-NN approach, which is different from a fixed *k*-nn approach; *v*-nn selects the neighbors based on a condition for each query compound. We used the median profile on the *v* training compounds that had a Tanimoto similarity greater than 0.70. We ignored any similar compound that appears in the external test set to avoid information leaks. Subsequently, we further discarded any compounds for which no feature profile was found directly or using the above *v*-nn approach. Thus, while the test sets for the DICTrank and DICTrank enriched datasets are the same, it is important to note that the training data for them vary for the models (as described in Supplementary Table S1) since we dropped compounds where no feature data could be found or matched.

For each of the eleven models, we used a Random Forest classifier, with hyperparameter optimization on the training data using a halving random search with a 5-fold stratified cross-validation with a random oversampling to account for class imbalance (as implemented in scikit-learn^39^). We used the best hyperparameter-optimized estimator and obtained out-of-fold predictions with a 5-fold stratified cross-validation. We used the out-of-fold predictions and the true labels to optimize the decision threshold for binary classification using the J statistic, calculated as the difference between the true positive rate and the false positive rate. This determines the threshold from ROC curve values where the J statistic is maximized. The model was finally refitted on the entire training dataset, and we used the optimized threshold to make final predictions based on the predicted probabilities of the external test set.

We trained two ensemble models to combine the models from the eleven feature spaces above. These were based on soft voting, which considered the mean of the scaled predicted probabilities of each mode (scaled according to the best threshold of each model). The first model considered only the six best-performing models (structural, physicochemical, MOA, CellScape, MOA with Cmax total, and CellScape with Cmax total) in the cross-validation (AUC>0.65). The second ensemble model considers all eleven models and thus is evaluated on the reduced external test set of 78 compounds where data from all feature spaces were available.

### Model evaluation and applicability domain

We evaluated the classifiers using the balanced accuracy, sensitivity (or recall), specificity, F1 score, Matthews Correlation Coefficient (MCC), AUC-ROC, and the AUC-PR, or precision-recall curve, which focuses on the positive class.

To evaluate the applicability domain of the models, for each compound in the external test set, we calculated the Tanimoto similarity of the nearest neighbor of the same DICTrank label (toxic/non-toxic) in the training dataset. We grouped compounds in 5 equal bins from Tanimoto similarity of 0.0 to 1.0 and evaluated the balanced accuracy and AUCPR in this range for the models used in this study.

### Statistics and Reproducibility

We have released the datasets used in this study which are publicly available at 10.6084/m9.figshare.24312274. We released the Python code for the models which are publicly available at https://github.com/srijitseal/DICTrank.

## Results and Discussion

In this study, we used various biological and chemical datasets to discern among the DICT concern categories, driving insights into the carefully annotated FDA DICTrank dataset. We also trained predictive models using these feature spaces. In particular, we used the Cell Painting data from Bray et al, which captures a wide array of cellular phenotypes after perturbation e.g. drug treatment, and has been shown to have a signal for various *in vitro* toxicity.^24, 49^ We also used experimental (from the Repurposing Hub^29^) and predicted bioactivity data derived from models trained on a mixture of publicly available and proprietary datasets (Ignota Labs CellScape^34^), mostly relating to inhibitory/antagonist mechanisms). For structure-derived feature spaces, we used Morgan fingerprints derived from chemical structures as well as physicochemical Mordred descriptors which are often related to pharmacokinetic properties (such as logD, molecular weight, solubility, permeability, etc.) and implicitly encode the bias between bioactivity classes and chemical structures.^50^ Finally, we looked at pharmacokinetic parameters for the peak unbound and total concentration of a drug molecule in plasma (Cmax).^51^ We organized and standardized various chemical and biological data, as shown in Table 2, to analyze their ability to predict DICTrank labels.

### DICTrank labels are highly concordant with SIDER labels

Among the 776 compounds present in both DICTrank and SIDER cardiac disorders datasets (Figure 2a), we found an 87.24% concordance rate in the annotations (labels) between the two datasets (Supplementary Table S3; SIDER labels have an F1 score of 0.91 when compared against DICTrank labels). This suggests that SIDER labels which ascertain cardiac disorder events reported as associated with each drug and are often dependent on aggregated dispersed public information and package inserts, agree with DICTrank labels which ascertain if a compound is classified as cardiotoxic by the FDA.

**Figure 2:**
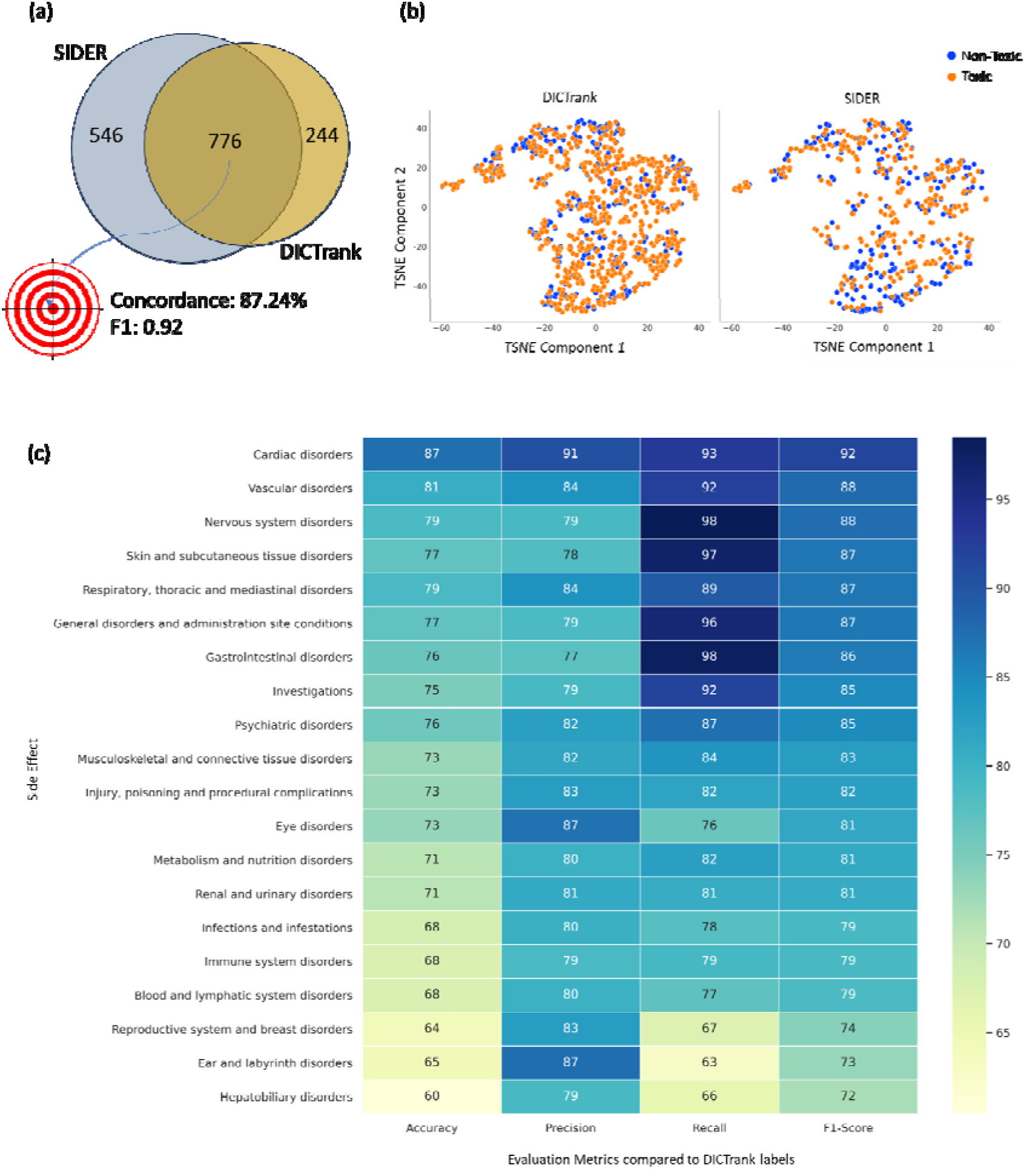
Comparison of the SIDER dataset with DICTrank: (a) the overlap and concordance (percentage of the total compounds with the same annotation) of DICTrank labels with SIDER cardiac disorder labels, (b) the overlay of SIDER and DICTrank chemical space in a principal component analysis using physicochemical properties, and (c) the positive predictive value of other side effects in SIDER for DICTrank labels (toxic/non-toxic).

The physicochemical space of SIDER and DICTrank generally overlap (Figure 2b), defined as a TSNE space for six physicochemical properties, namely, molecular weight, topological polar surface area, number of rotatable bonds, hydrogen bond donors and acceptor, and the computed logarithm of the partition coefficient. Still, compounds exclusively available in the SIDER dataset could help enrich nontoxic compounds in areas of the chemical space where DICTrank only covers toxic compounds. We see a similar trend for a chemical space defined in fragment fingerprints space from DataWarrior^40^ (Supplementary Figure S1). Therefore, we chose to assess whether adding SIDER compounds to DICTrank compounds improved predictive ability. Interestingly, other categories of SIDER adverse effects were highly correlated to DICTrank (Figure 2c); the interrelationships of vascular disorders and nervous system disorders are well known.^5, 54^ Overall, drug adverse events, as recorded in SIDER, have a high concordance with DICTrank labels from the FDA and there is a strong rationale to rely on both resources.

### Maximum total and unbound compound concentration in plasma predict cardiotoxicity

We next determined if a high Cmax indicated compounds more likely to be cardiotoxic as seen in the case of doxorubicin where cardiotoxicity was found to be Cmax dependent.^55^ As a single parameter, Cmax was not sufficiently discerning to differentiate between compounds that fall under the ’most-concern’ and ’less-concern’ categories as per the DICT concern classification (Figure 3). However, for both peak total plasma levels and peak unbound (active) plasma levels’ Cmax, the median distributions were significantly distinguishable between cardiotoxic and non-toxic compounds (Figure 3) suggesting that Cmax can be a useful parameter in determining cardiotoxicity.

**Figure 3:**
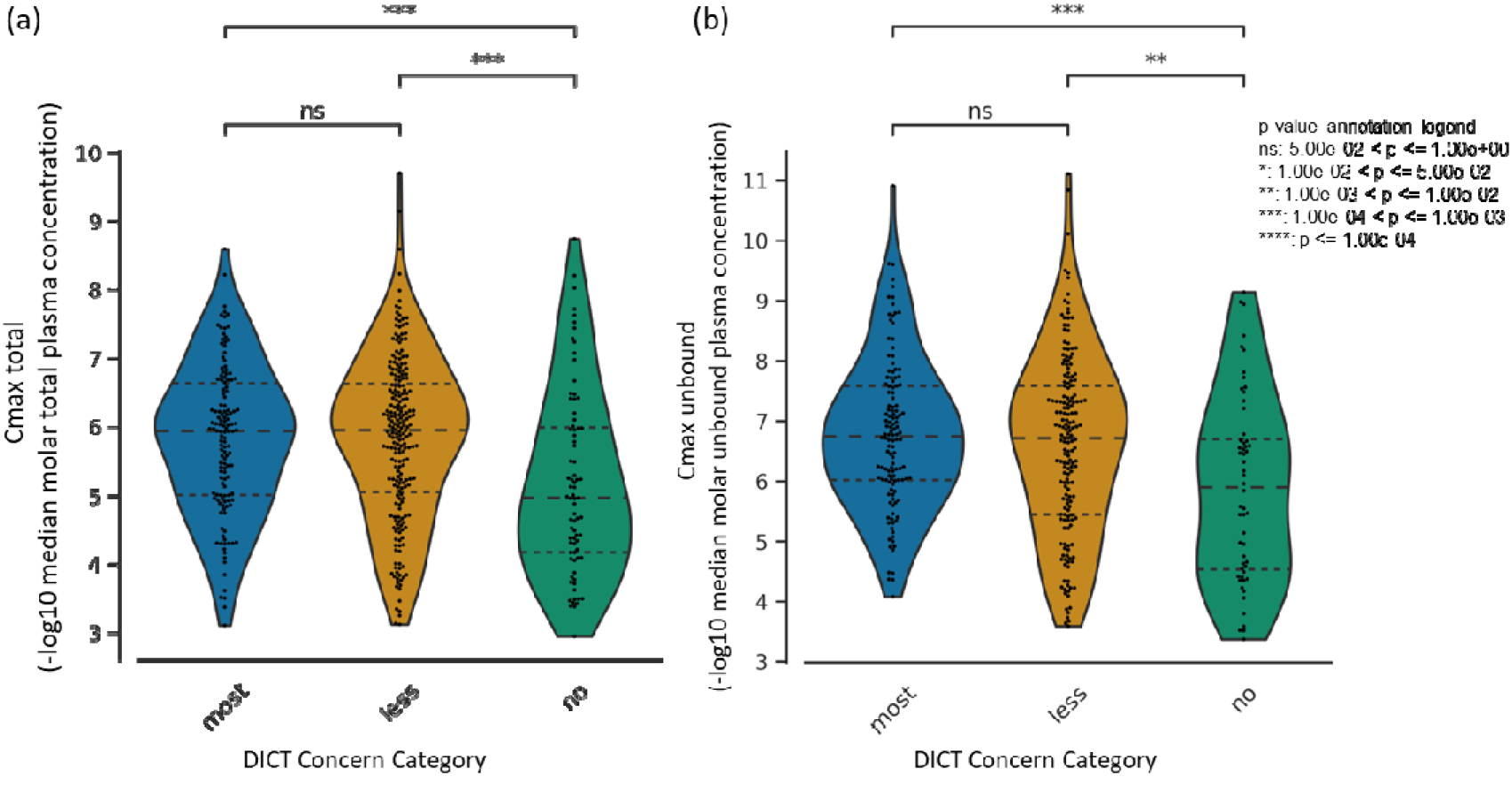
The distribution of (a) peak total concentration in plasma and (b) peak unbound (active) concentration in plasma for each drug in the DICTrank dataset across the three DICT concern categories.

### Cyclooxygenase inhibition is predictive of cardiotoxicity concern

Turning to manual annotations of compound mechanisms of action and/or targets, we found that cyclooxygenase inhibitors^56^ along with tyrosine kinase receptor inhibitors were the most significant annotations differentiating the various DICT concern categories (Table 3); this is plausible given cyclooxygenase inhibition, besides reducing inflammation, can also lead to increased blood pressure^57^ while tyrosine kinase receptor inhibition can induce endoplasmic reticulum stress and inflammation in cardiomyocytes.^58^ In agreement with this, known targets of prostaglandin endoperoxide synthases (PTGS1 and PTGS2 genes, which encode cyclooxygenases COX-1 and COX-2) could significantly distinguish among most-, less- and no-DICT concern categories (Table 3).

**Table 3:**
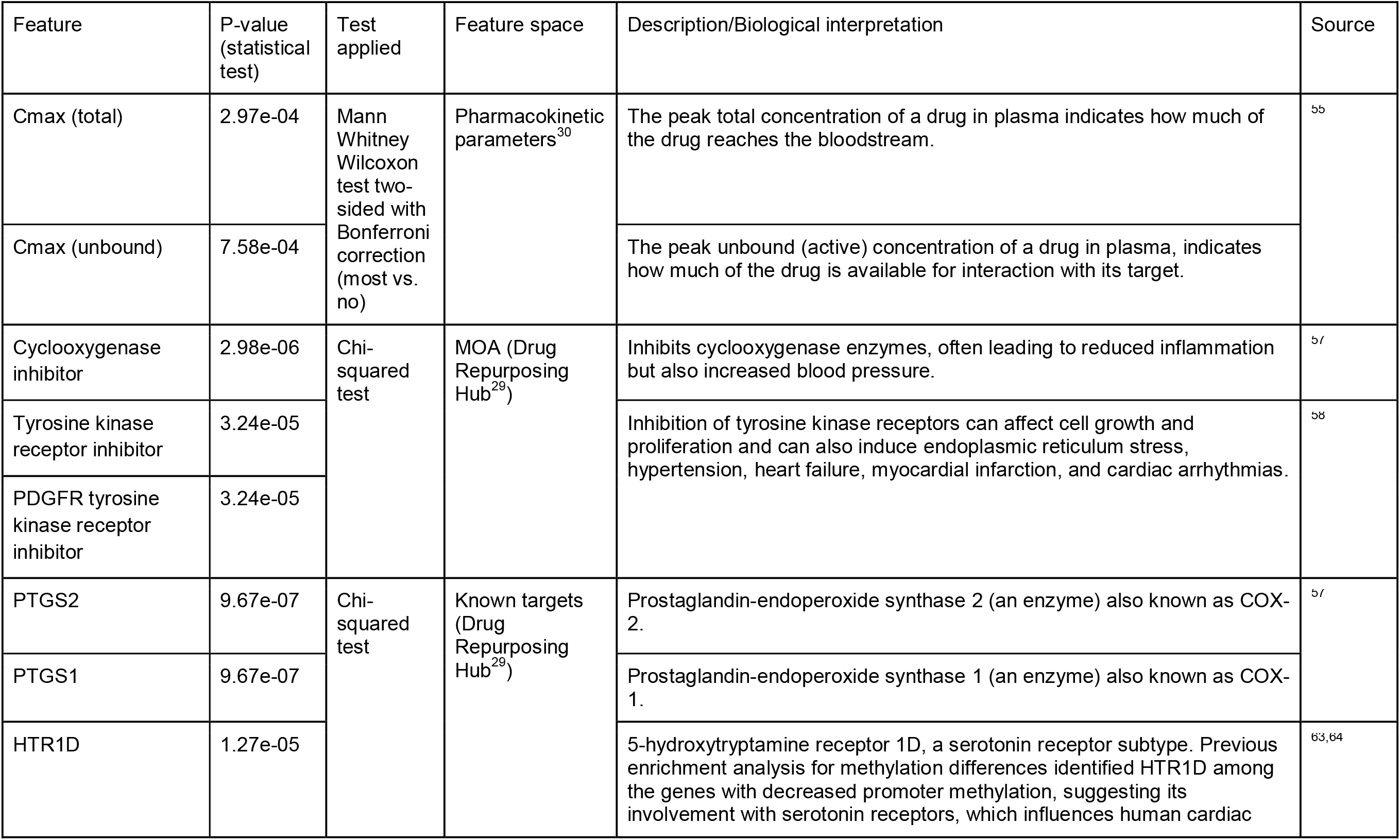
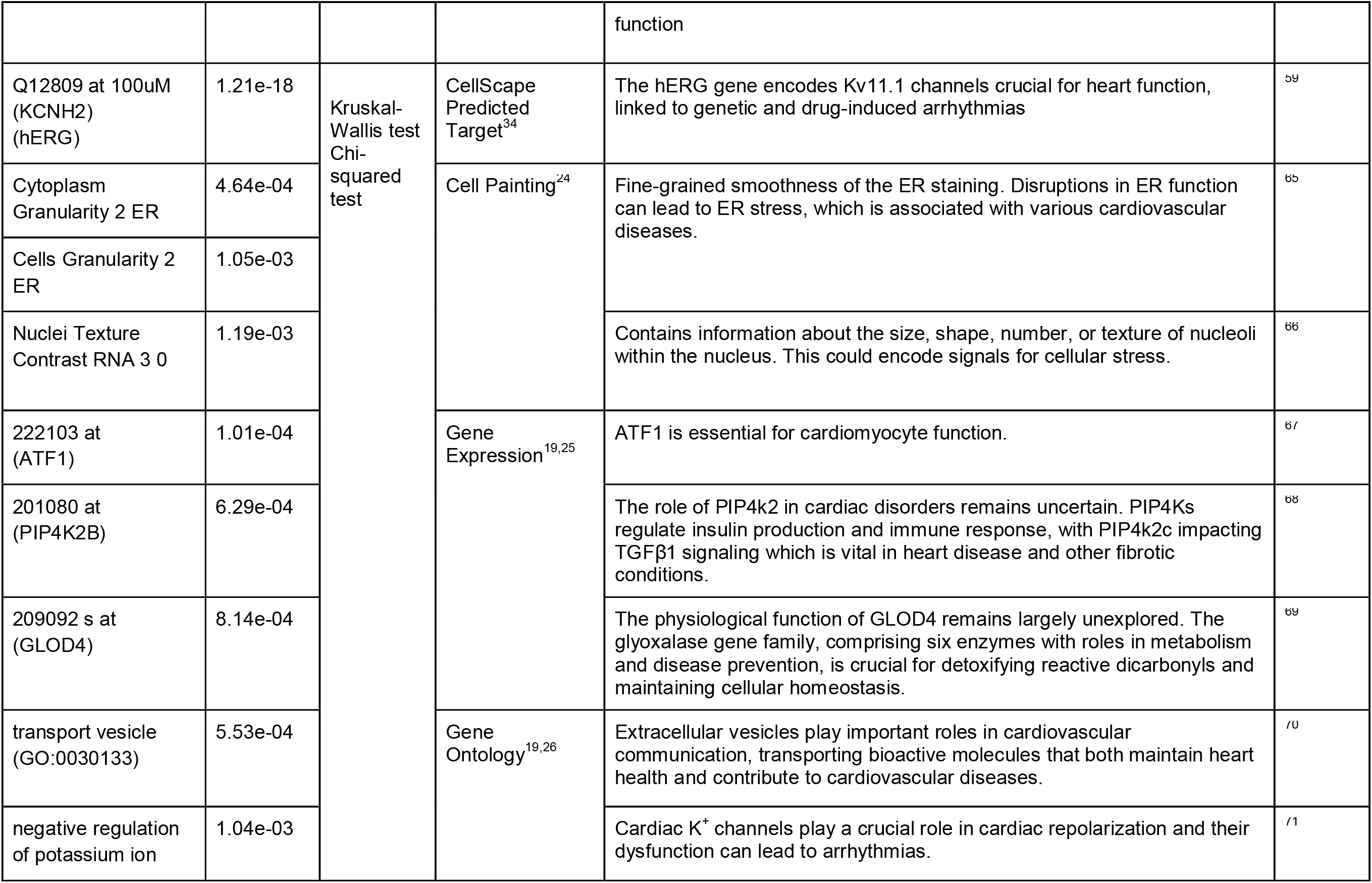
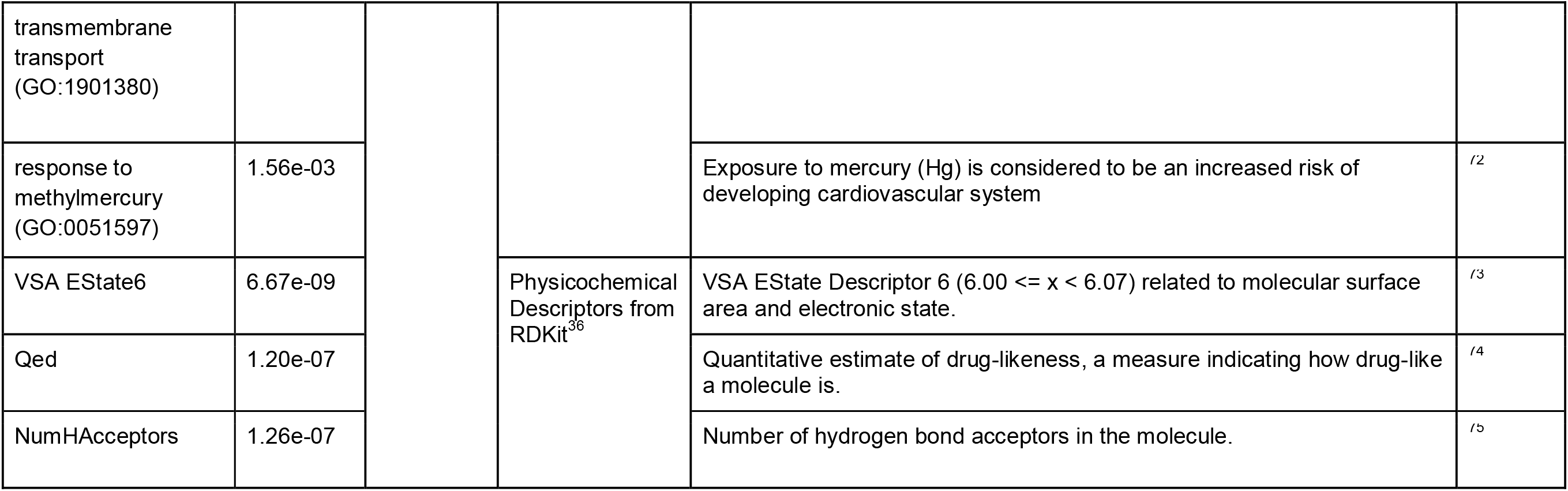
Features from chemical and biological data sources that have a particularly high significance in the difference of distributions for the three DICT Concern Categories (Most, Less, No-concern).

### CellScape-predicted protein targets such as hERG are predictive of cardiotoxicity

Among CellScape-predicted protein targets, the predicted activity of compounds against KCNH2 is best differentiated among the three DICT concern categories. The KCNH2 gene, also known as the human ether-à-go-go-related gene (hERG), is well known for its significance in the cardiac electrical cycle and hERG inhibition can lead to cardiac arrhythmias.^59^ We also found that the top three features to distinguish the two DICTrank labels (cardiotoxic versus non-toxic) were α-l-fucosidase I, P-selectin, and carbonic anhydrase IX. The activity of plasma α-l-fucosidase has been pinpointed as a potential biomarker for cardiac hypertrophy and complements the currently used marker, atrial natriuretic peptide.^60^ Elevated amounts of soluble P-selectin in the blood are evident in various heart-related conditions, like coronary artery disease, hypertension, and atrial fibrillation.^61^ Carbonic anhydrase IX plays a role in managing the intracellular pH in the heart muscle, a vital for the heart’s functionality.^62^

### Hypothesis-free omics data for cardiotoxicity are related to mechanisms of action

Omics data sources such as Cell Painting (imaging), gene expression, and Gene Ontology features cover a broad swath of biology, not specifically targeted to cardiac function. For Cell Painting, the fine-grained smoothness of the ER in the cytoplasm and RNA in the nucleus were the top features that differed significantly among toxicity classes. This is plausible given disruptions in ER function can lead to ER stress, which is associated with various cardiovascular diseases.^65^ For the gene expression feature space, activating transcription factor 1 (ATF1), which is essential for cardiomyocyte function, was the top feature. The other two gene expression features that could distinguish DICT concern categories were phosphatidylinositol-5-phosphate 4-kinase type 2 beta (PIP4K2B) and glyoxalase domain containing 4 (GLOD4); both have indirect links to heart disease and other fibrotic conditions (Table 3). Among Gene Ontology annotations, we found that biological processes related to vesicle transport, potassium ion transmembrane transport, and response to methylmercury could best differentiate signals for concern categories. This is plausible given cardiomyocytes rely on vesicular transport for various functions, including the delivery of membrane proteins and lipids. The potassium ion channels play crucial roles in cardiac cell electrical activity and dysregulation can lead to arrhythmias and other heart complications.^70, 71^ Exposure to mercury (Hg) is also considered a risk for ischemic heart disease.^72^

### Physicochemical Properties can differ among DICT concern categories

Among the various molecular descriptors evaluated in our study, VSA_EState6 could significantly distinguish among the DICT-concern categories. This electrotopological state descriptor aggregates the differences in electronegativity between an atom and its neighboring atoms in a molecule, adjusted by their relative distances while focusing on atoms with specific van der Waals surface area.^73^ This suggests that specific electronic and spatial properties are captured by the VSA_EState6 descriptor, although difficult to interpret directly. The second predictive feature, Qed, captures a quantitative estimation of the drug-likeness score that encapsulates the underlying distribution data for a range of drug properties.^74^ The third predictive feature, NumHAcceptors refers to the number of hydrogen bond acceptors in the compound. Munawar et al. showed that the most potent hERG inhibitors typically possess two aromatic groups, one hydrophobic group, and one hydrogen bond acceptor, at specific relative distances from each other.^75^

### Structural alerts from DICTrank can detect compounds causing cardiac disorders from a retrospective analysis of DrugBank

We determined 59 structural alerts that distinguish cardiotoxic and non-toxic compounds in the DICTrank dataset (Figure 4). Two structural alerts had a high positive predictive value (PPV) for the DICT most-concern category, including one with aromatic rings. Aromatic rings can lead to π-stacking or hydrophobic interactions with aromatic rings of amino acids within the hERG channel cavity increasing the potential for blocking and subsequent cardiotoxic effects.^76^ Six structural alerts distinguished toxic versus non-toxic compounds with a positive predictive value of 1 and more than ten occurrences in the dataset (the PPV was used to filter the structural alerts, hence is not an evaluation metric here). Structural alerts with tertiary amines were consistently protonated at physiological pH in the DICTrank dataset, suggesting their importance in biological activity and hERG channel binding.^77, 78^ It is also known that compounds with secondary amine (more hydrogen bond donor number) are likely to be less potent hERG inhibitors compared to tertiary amine (less hydrogen bond donor number).^78^

**Figure 4:**
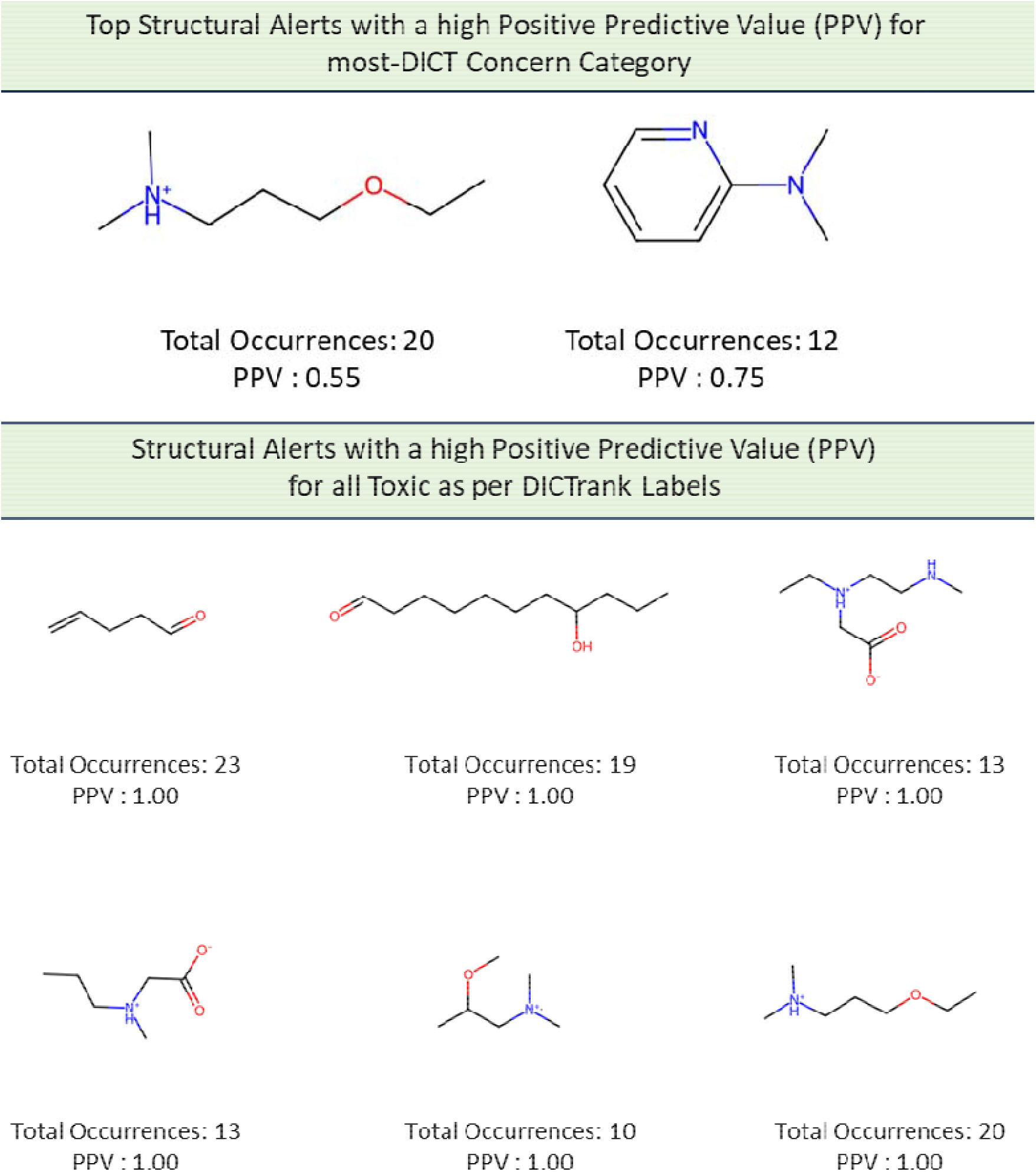
Structural alerts for (top) the most-concern DICT category and (bottom) DICTrank labels with more than ten occurrences and a PPV>0.6 for compounds in the DICTrank dataset.

We next analyzed compounds in DrugBank^47^ for the presence of at least one of the two structural alerts above for the most-concern category. We annotated these hits with heart-related side effects from SIDER^32^ and their current status (approved, withdrawn, etc.) as indicated in DrugBank. We found six approved drugs, some experimental/investigational, with reported cardiac disorders from SIDER (Table 4). These compounds spanned different classes of compounds, with the presence of a tertiary amine that remains protonated or aminopyridine rings as defined by the structural alerts. We found evidence in the literature for the risk of cardiovascular disorders for three of the six compounds, namely, ipratropium, tiotropium, and mivacurium.^79–81^ Overall, our analysis shows that the DICTrank dataset is a rich source of cardiotoxicity-causing compounds, with the potential to be used to build pharmacophore models and evaluate compounds with reported adverse events for their potential mechanisms of toxicity. Overall, we could detect multiple approved drugs that match the structural alerts for both the DICT most-concern category (as shown in Table 4) and for DICTrank labels for cardiotoxicity (further details in Supplementary Figure S2).

**Table 4:**
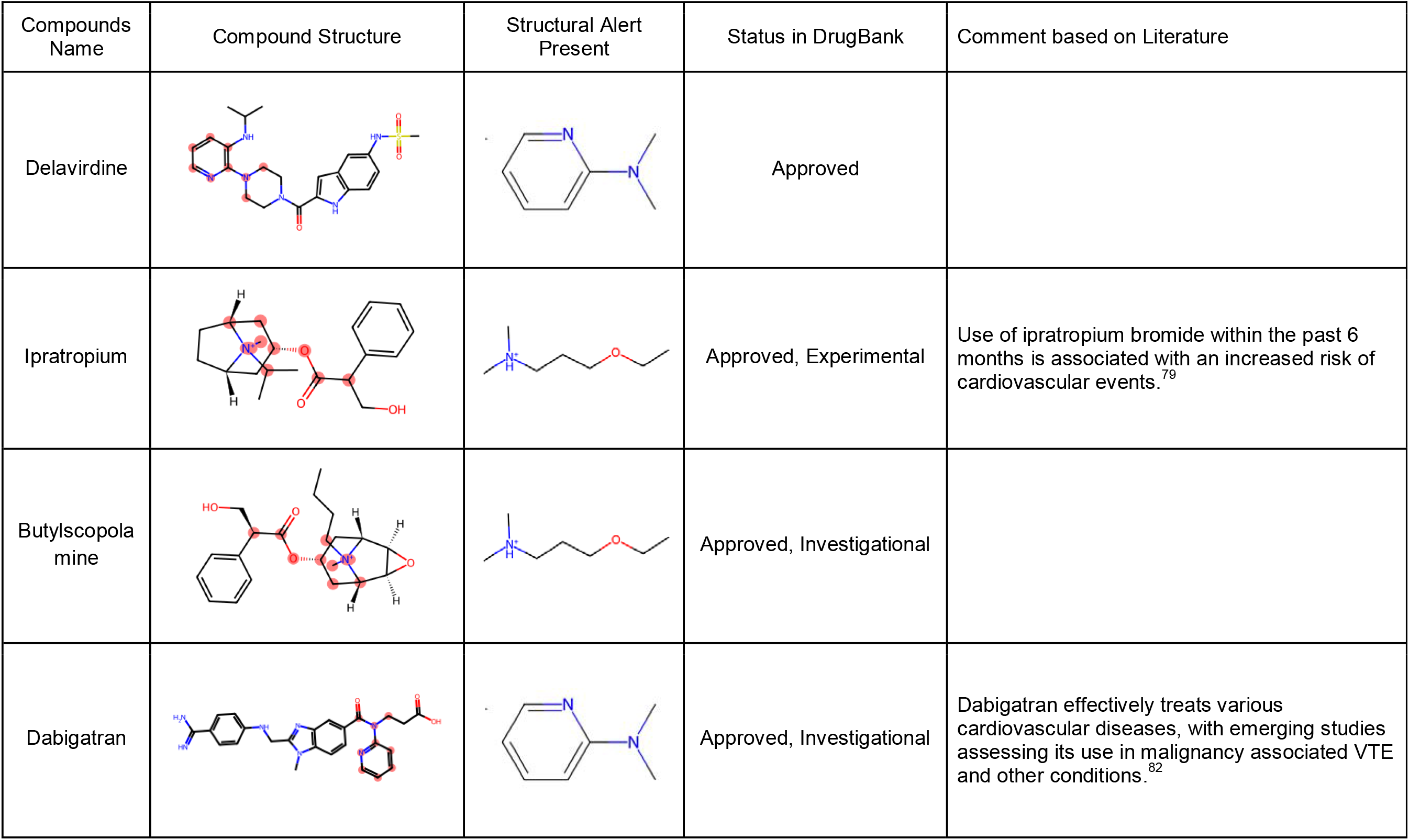
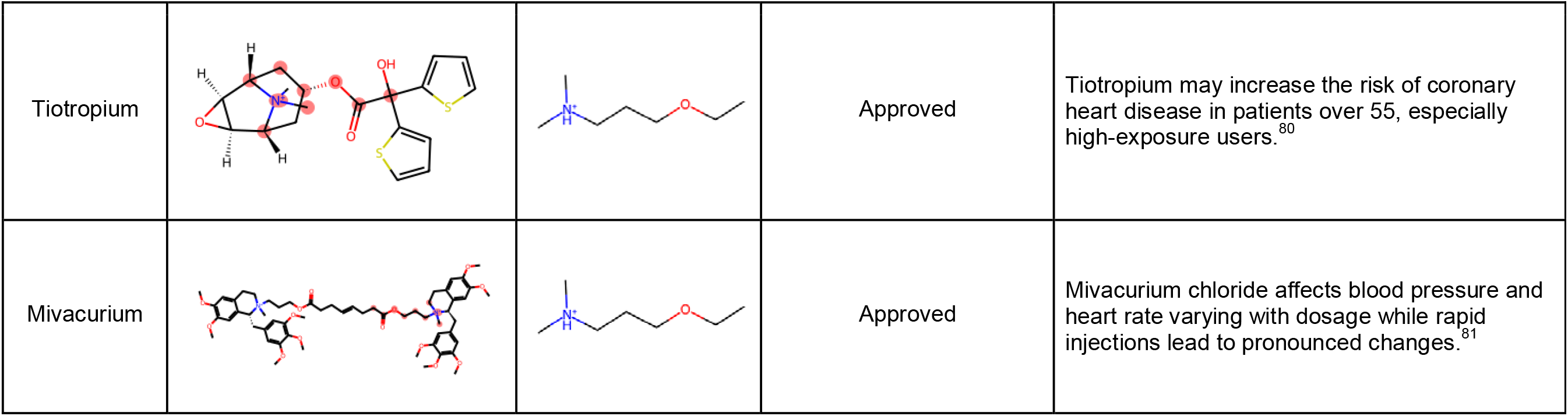
Six hits from SIDER with the structural alerts for the DICT most-concern category.

| Compounds Name | Compound Structure | Structural Alert Present | Status in DrugBank | Comment based on Literature |
| --- | --- | --- | --- | --- |
| Delavirdine |  |  | Approved |  |
| Ipratropium |  |  | Approved, Experimental | Use of ipratropium bromide within the past 6 months is associated with an increased risk of cardiovascular events.<sup>79</sup> |
| Butylscopolamine |  |  | Approved, Investigational |  |
| Dabigatran |  |  | Approved, Investigational | Dabigatran effectively treats various cardiovascular diseases, with emerging studies assessing its use in malignancy associated VTE and other conditions.<sup>82</sup> |
| Tiotropium |  |  | Approved | Tiotropium may increase the risk of coronary heart disease in patients over 55, especially high-exposure users.<sup>80</sup> |
| Mivacurium |  |  | Approved | Mivacurium chloride affects blood pressure and heart rate varying with dosage while rapid injections lead to pronounced changes.<sup>81</sup> |

### Predictive models for DICTrank labels

Finally, given the promising signals seen in each data type as described above, we evaluated whether cardiotoxicity might be predicted using the data sources currently publicly available. Several data sources contained sufficient information to successfully train models to predict DICTrank labels (Table 2). We trained 11 models on four types of training data: the DICTrank compounds alone and DICTrank compounds enriched with cardiotoxic/non-toxic/all compounds in the SIDER dataset (as shown in Supplementary Table S1). A direct comparison of the predictive value of data sources is not possible due to the incomplete intersection of compounds with available data of each type. Still, we fixed the held-out test set of compounds to be those where data was available for all feature spaces such that only the training set of compounds varied among data sources. We trained two ensemble models, one using six models (structural, physicochemical, MOA, CellScape, MOA with Cmax total, and CellScape with Cmax total) that performed relatively well on the internal cross-validation (evaluation metrics from cross-validation for all feature space and dataset combinations, are given in Supplementary Table S4). This ensemble model was evaluated on an external test set of 90 compounds. Another ensemble model was built on all eleven models, which required testing on a smaller held-out test set due to the limited overlap of data. Evaluation metrics for all models are given in Supplementary Table S5.

Looking at each data source independently, we found that models using Mordred descriptors evaluated on the 90 compounds held-out test set (AUC: 0.84, AUCPR: 0.93; random AUC: 0.50, AUCPR: 0.72) performed better compared to models trained on predicted protein targets (AUC: 0.77, AUCPR: 0.89) and MOA annotations with Cmax (total) (AUC: 0.77, AUCPR: 0.90) (Figure 5a, b). In fact, models using Mordred descriptors were as good as the ensemble of six selected models (AUC: 0.83, AUCPR: 0.92; random AUCPR: 0.72) also evaluated on the 90 compounds held-out test set (Supplementary Figure S3). Further, models across most datasets performed with high AUCPR and F1 scores, with top-performing models using Mordred descriptors (AUCPR: 0.93; random AUCPR: 0.72) and ensemble models (AUCPR: 0.93 for both ensemble models) when using the DICTrank dataset directly (Supplementary Figures S3a and S3b). Exceptions were models using the broad-based omics data -Cell Painting, Gene Expression, and Gene Ontology - where the performance was relatively poor and similar to random predictions according to the distribution of respective training data. This lack of predictive power may be inherent to the data sources but could also be due to the highly unbalanced and sparse training data available for these data sources (see Supplementary Table S2). When comparing the models evaluated with the smaller test set (Supplementary Figure S3), we found that models trained on the DICTrank dataset enriched with all SIDER compounds and using MOA data with Cmax (unbound) (AUCPR: 0.93, random AUCPR: 0.73) performed equally as the ensemble models that used predictions from all eleven models trained on just the DICTrank dataset (AUCPR: 0.93; random AUCPR: 0.73). Overall, a strong detection of cardiotoxicity was seen equally among the ensemble model and models using physicochemical descriptors.

**Figure 5:**
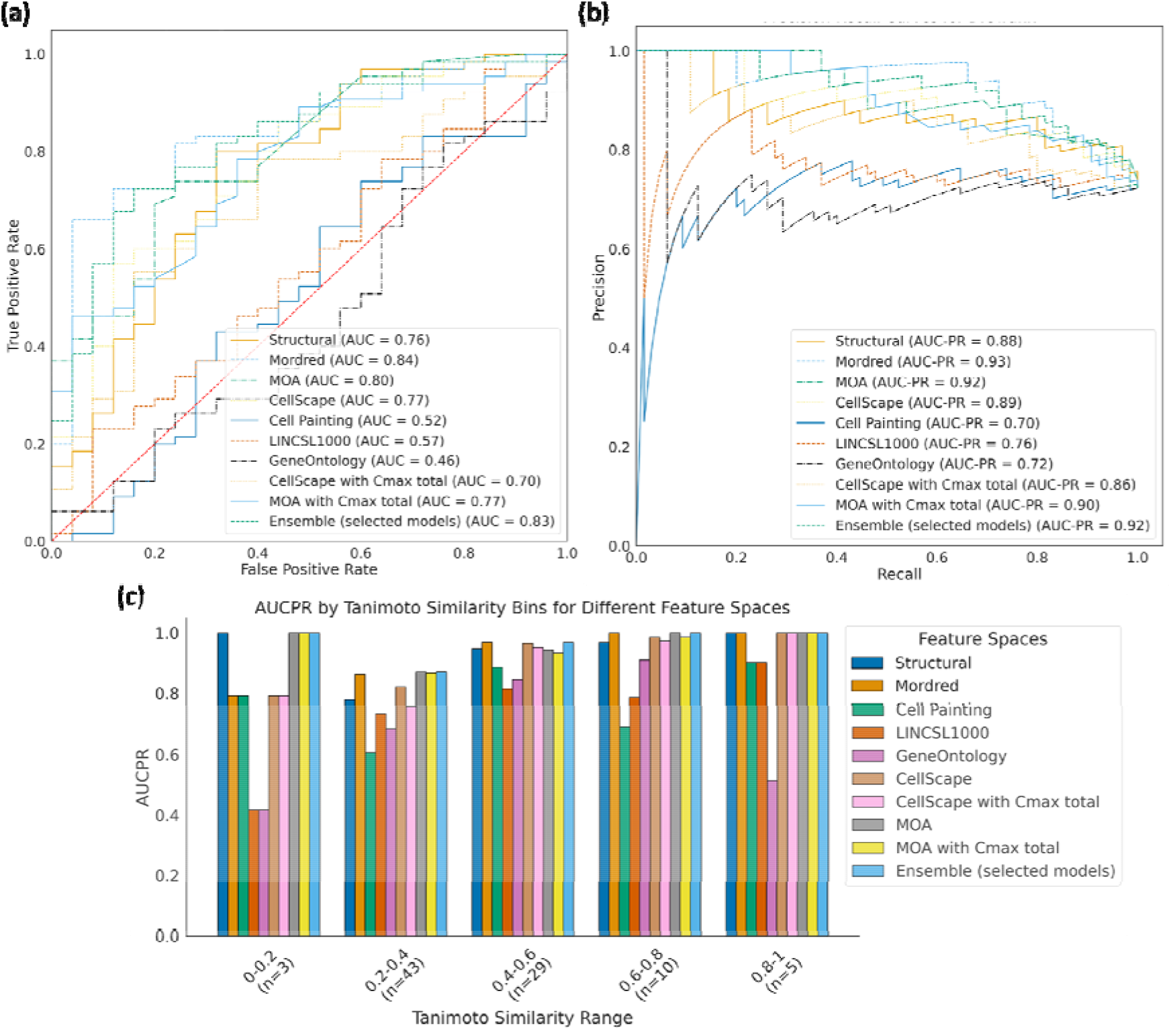
Comparison of evaluation metrics models built in this study with an external test set of 90 compounds evaluated by the (a) AUC-ROC and (b) AUCPR and (c) the performance of each model across compounds that are similar (dissimilar) to the training data. The ensemble mode in (c) is based on the models built on six data sources listed in the main text and evaluation is for 90 held-out compounds; results for the ensemble using 11 data sources and 78 held-out compounds are in Supplementary Figure S3.

We next analyzed the applicability domain of these models based on evaluating the quality of prediction for groups of compounds that are structurally dissimilar to the training data. We found that ensemble models and models using MOA annotations perform consistently well across the similarity range (Figure 5c). Models using Mordred descriptors, on the other hand, perform with slightly lower AUC-PR when compounds are structurally dissimilar to training data.

Finally, we predicted the DICTrank labels for 82 unique compounds that were labeled ambiguous in the original DICTrank dataset (Supplementary Table S6). We used Mordred descriptors and retrained the model on all 1020 compounds (training and held-out compounds) in DICTrank, except for the ambiguous compounds. We found that 43 of the 82 compounds were predicted to be cardiotoxic and 39 were predicted to be non-toxic and provided this list to the community for further study (Supplementary Data S6).

### Limitations of this Study

While we considered in this study various chemical and biological data sources, it is important to remember that conclusions are based on limited data. Certain feature spaces contain features that are computed based on chemical structure, such as CellScape target predictions and physicochemical properties, while datasets such as MOA and SIDER are manually gathered and have evidence of the presence and absence of evidence annotations. To train models using feature spaces such as Cell Painting, Gene Expression, and Gene Ontology datasets, we dropped compounds where we could not find profiles (whether experimentally captured or imputed based on matching to highly similar compound profiles using the *v*-nn approach). The amount of training data (and also the class balance of SIDER/DICTrank labels) is lower for these models. Although we compare data sources using the same test compounds, the varying amounts of training data, and the differing types of compounds represented therein, can disadvantage some data sources versus others, such that we cannot with certainty compare the signal contained across the feature spaces. The poor performance of -omics data should therefore not yet be attributed to representing the signal in the feature space. Rather in this study, we aim to evaluate the signal present in the data that is available and build the best predictive models possible with public data. In the future, the availability of more data, for example, Cell Painting from JUMP-CP^83^ and Recursion RxRx3^84^ will significantly improve our ability to ascertain the presence of a signal for cardiotoxicity in -omics data.

### Conclusions

In this work, we used biological data and chemical data (Figure 1) to predict drug-induced cardiotoxicity. We determined the feature contained in each data source that most differed between the most-concern versus non-toxic category for DICTrank and found these could drive mechanistic insights. Features from data sources such as predicted protein targets and annotated MOAs that could distinguish the DICT concern categories resembled activity against targets (ion channels in particular) that are mechanistically most plausible. We further evaluated these feature spaces using machine learning to build the first predictive models of DICTrank. Our findings indicate that models relying on physicochemical properties trained on larger training datasets performed on par with the ensemble models based on diverse data sources. The exploratory data analysis in this study suggests that as more - omics data becomes accessible in the future, it will enhance our ability to predict cardiotoxicity. Therefore, for the present, when constructing models using public datasets, we advocate the use of Mordred descriptors and predicted targets (based on chemical structure), since these computed properties are readily available for compounds; they do not require experimental data and could be used to build models for cardiotoxicity. In the future, using biological data we can look into the biological pathways and mechanisms of DICT leading to better drug design and safer therapeutic strategies.

### Associated Content

Supplemental Information. Supporting Information is available. Supporting Information (PDF). We released the Python code for our models which are publicly available at https://github.com/srijitseal/DICTrank/ and all datasets at https://doi.org/10.6084/m9.figshare.24312274.v1

## Acknowledgments

This work was performed using resources provided by the Cambridge Service for Data-Driven Discovery (CSD3) operated by the University of Cambridge Research Computing Service (www.csd3.cam.ac.uk), provided by Dell EMC and Intel using Tier-2 funding from the Engineering and Physical Sciences Research Council (capital grant EP/T022159/1), and DiRAC funding from the Science and Technology Facilities Council (www.dirac.ac.uk). Cartoons in TOC Figure and Figure 1 were created with DALLEv2 (https://openai.com/dall-e-2), Microsoft Designer (https://designer.microsoft.com/) and Bioicons (https://bioicons.com) which compiled images from the Database Centre for the life sciences/ TogoTV (https://togotv.dbcls.jp) and Servier (https://smart.servier.com).

## Author Declarations

S Singh and AEC serve as scientific advisors for companies that use image-based profiling and Cell Painting (AEC: Recursion, SyzOnc, S Singh: Waypoint Bio, Dewpoint Therapeutics) and receive honoraria for occasional talks at pharmaceutical and biotechnology companies. OS declares shares in Phenaros Pharmaceuticals. LGH is an employee at Ignota Labs where CellScape is a proprietary software. All other authors declare no relevant competing interests.

## Author contributions

All authors have approved the final version of the manuscript. S Seal designed and performed exploratory data analysis, and implemented, and trained the models. LHG trained CellScape for the prediction of drug targets. S Seal, AEC, and S Singh analyzed the biological interpretation of morphological features. S Seal, OS, and AB analyzed the performance of machine learning models. S Seal wrote the manuscript with extensive discussions with all authors. All the authors (S Seal, OS, LHG, MGO, AEC, AB, and S Singh) reviewed, edited, and contributed to discussions on the manuscript.

