## Supplementary Figures for "Insights into Drug Cardiotoxicity from Biological and Chemical Data: The First Public Classifiers for FDA DICTrank"

Documentation: <https://broad.io/DICTrank_Predictor>

Code: <https://github.com/srijitseal/DICTrank>

Datasets: 10.6084/m9.figshare.24312274

*Correspondence:

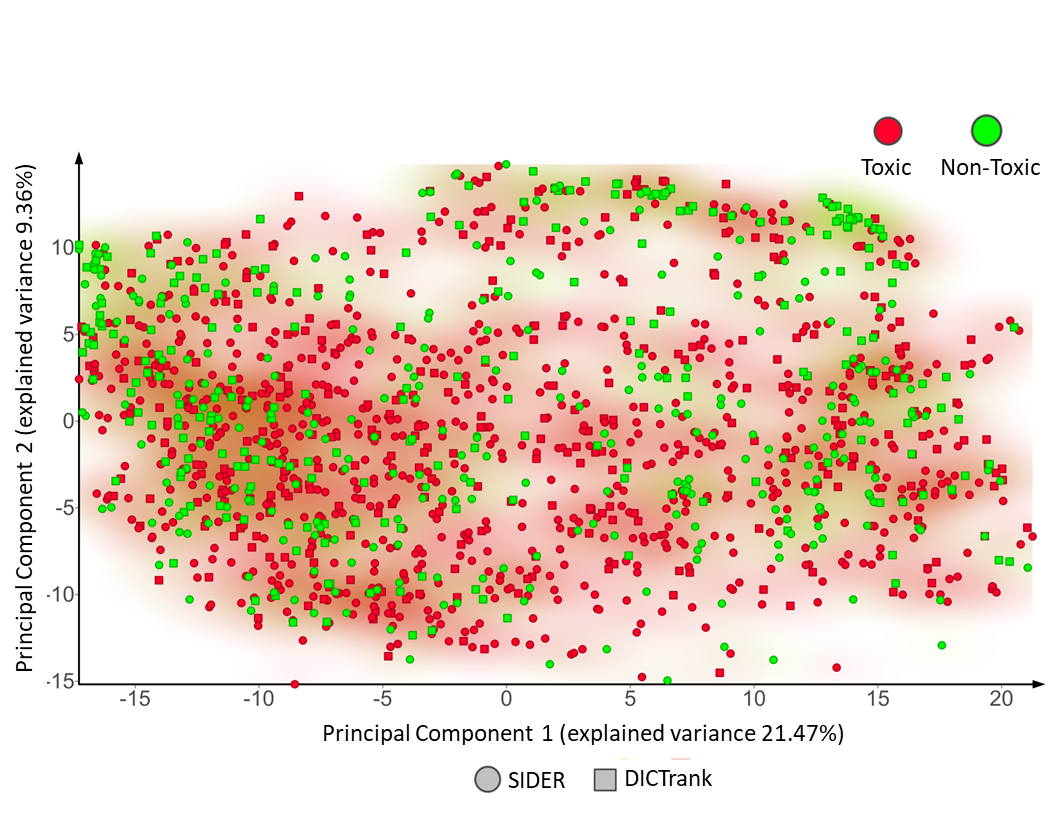

Figure S1: The chemical space of SIDER compounds and DICTrank compounds as defined by FragFP fingerprints (chemical fingerprints) in a principal component analysis. SIDER and DICTrank compounds fairly overlap (with the PCA explained variance of 21.47%).

**Figure S2:** Hits from DrugBank compounds containing structural alerts from the top 6 presented in Figure 4(b).

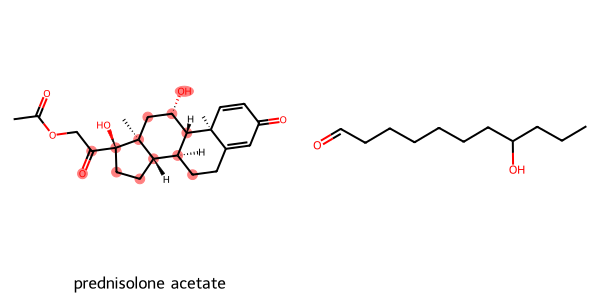

['Cardiac disorders']

Additional Data: ['approved']

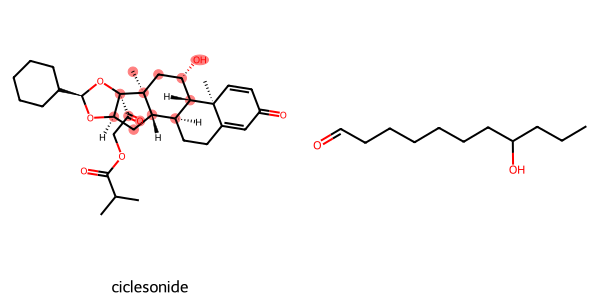

['Cardiac disorders']

Additional Data: ['approved', 'investigational']

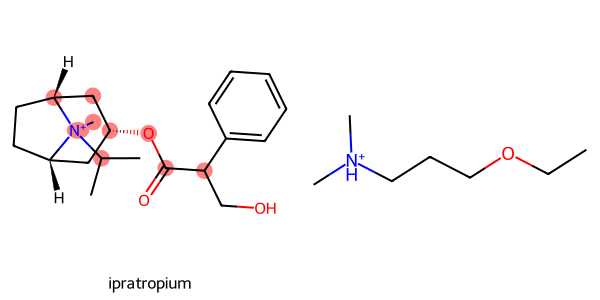

['Cardiac disorders']

Additional Data: ['approved', 'experimental']

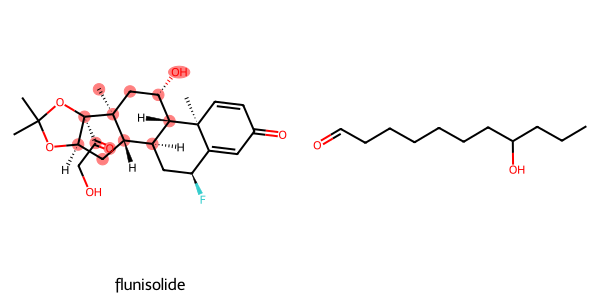

['Cardiac disorders']

Additional Data: ['approved', 'investigational']

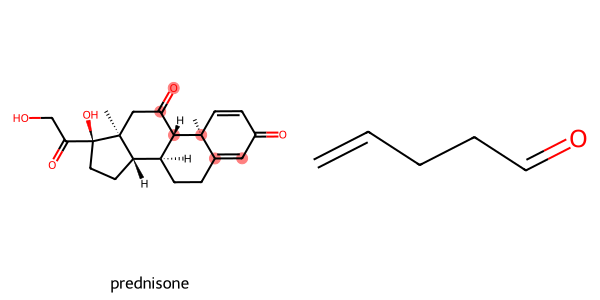

['Cardiac disorders']

Additional Data: ['approved']

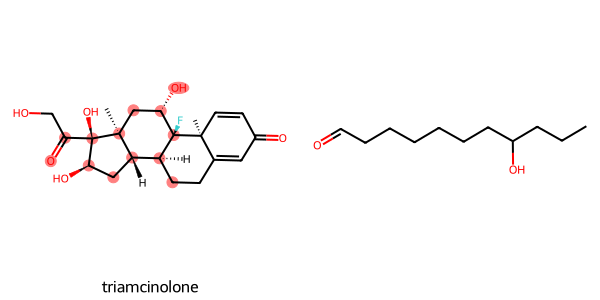

['Cardiac disorders']

Additional Data: ['approved']

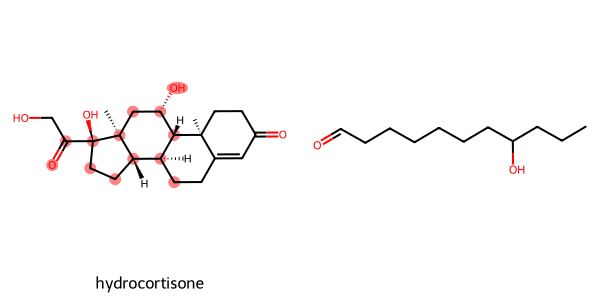

['Cardiac disorders']

Additional Data: ['approved']

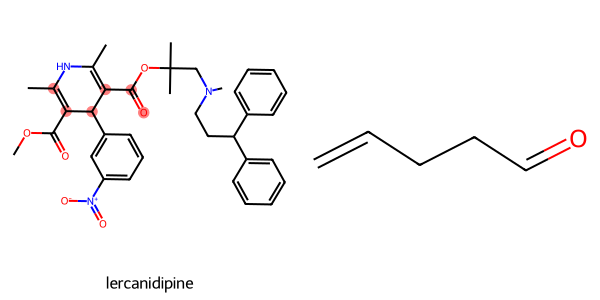

['Cardiac disorders']

Additional Data: ['approved', 'investigational']

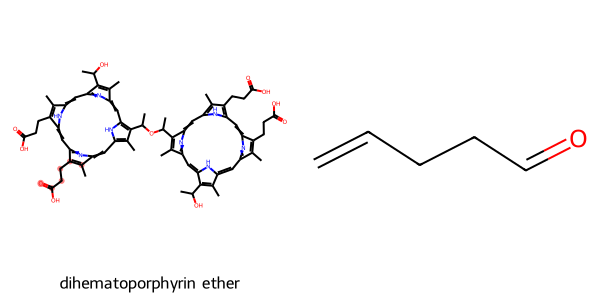

['Cardiac disorders']

Additional Data: ['investigational']

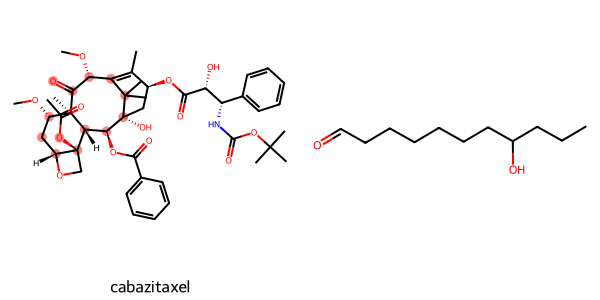

['Cardiac disorders']

Additional Data: ['approved']

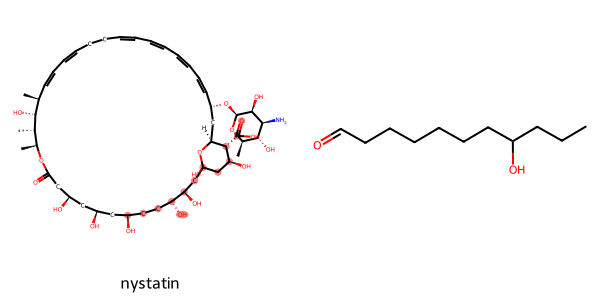

['Cardiac disorders']

Additional Data: ['approved']

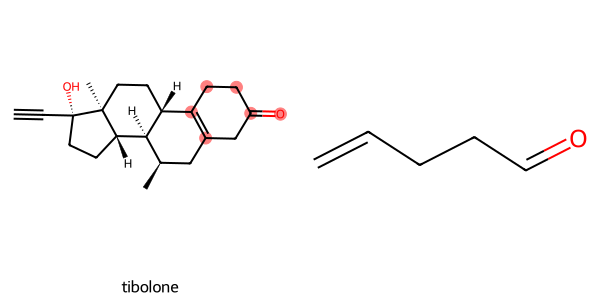

['Cardiac disorders']

Additional Data: ['approved', 'investigational']

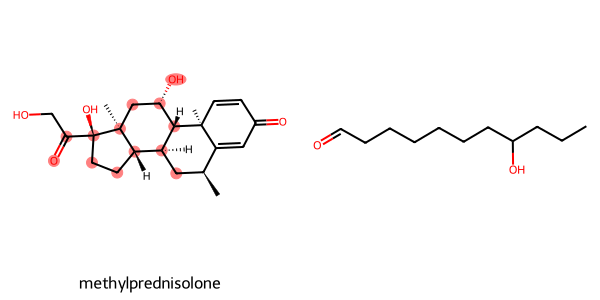

['Cardiac disorders']

Additional Data: ['approved']

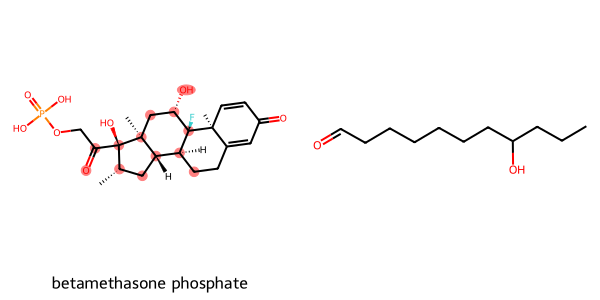

['Cardiac disorders']

Additional Data: ['approved']

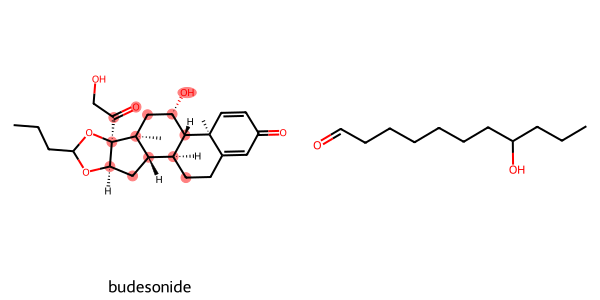

['Cardiac disorders']

Additional Data: ['approved']

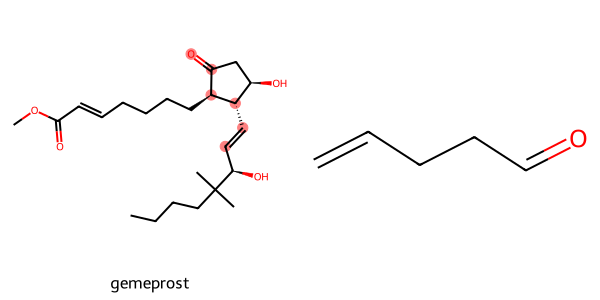

['Cardiac disorders']

Additional Data: ['approved', 'withdrawn']

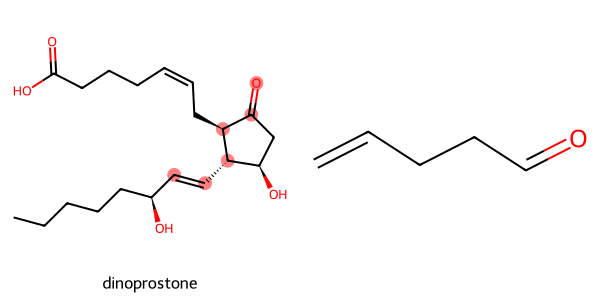

['Cardiac disorders']

Additional Data: ['approved']

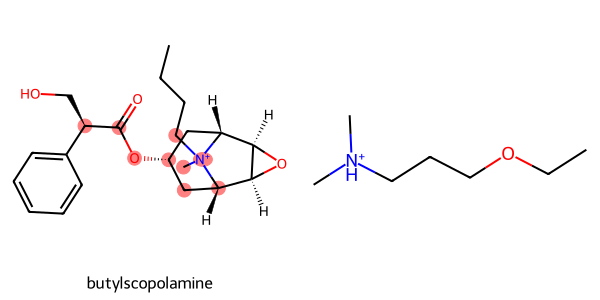

['Cardiac disorders']

Additional Data: ['approved', 'investigational']

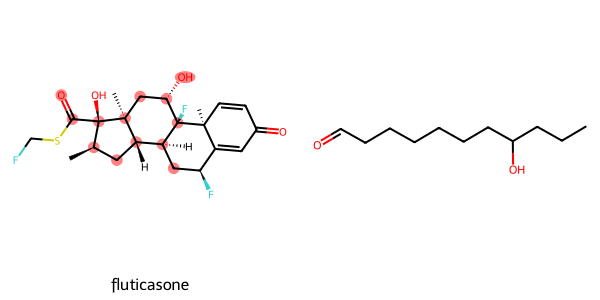

['Cardiac disorders']

Additional Data: ['approved', 'experimental']

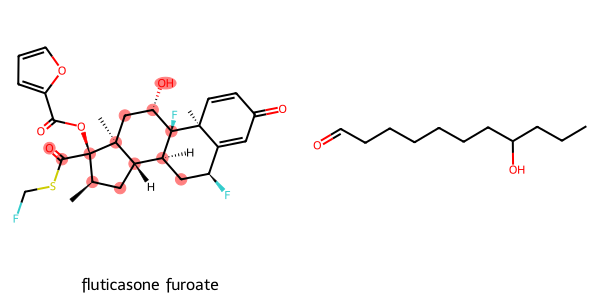

['Cardiac disorders']

Additional Data: ['approved']

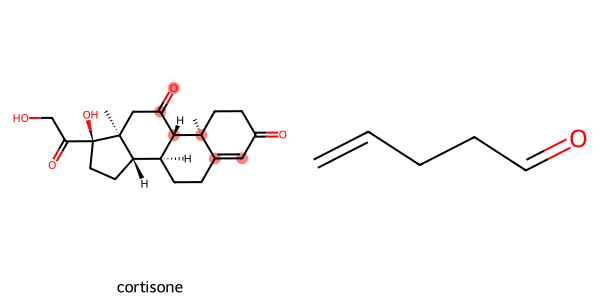

['Cardiac disorders']

Additional Data: ['experimental']

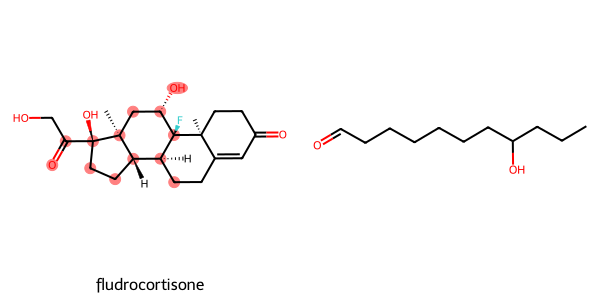

['Cardiac disorders']

Additional Data: ['approved', 'investigational']

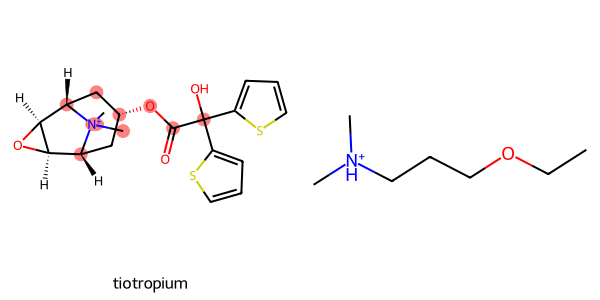

['Cardiac disorders']

Additional Data: ['approved']

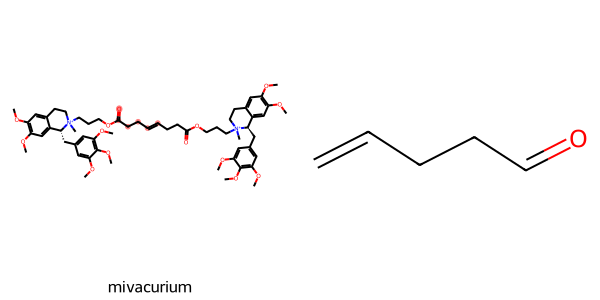

['Cardiac disorders']

Additional Data: ['approved']

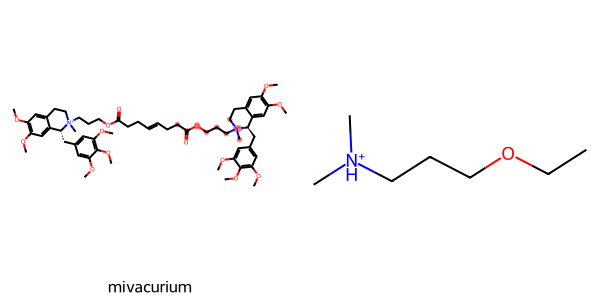

['Cardiac disorders']

Additional Data

: ['approved']

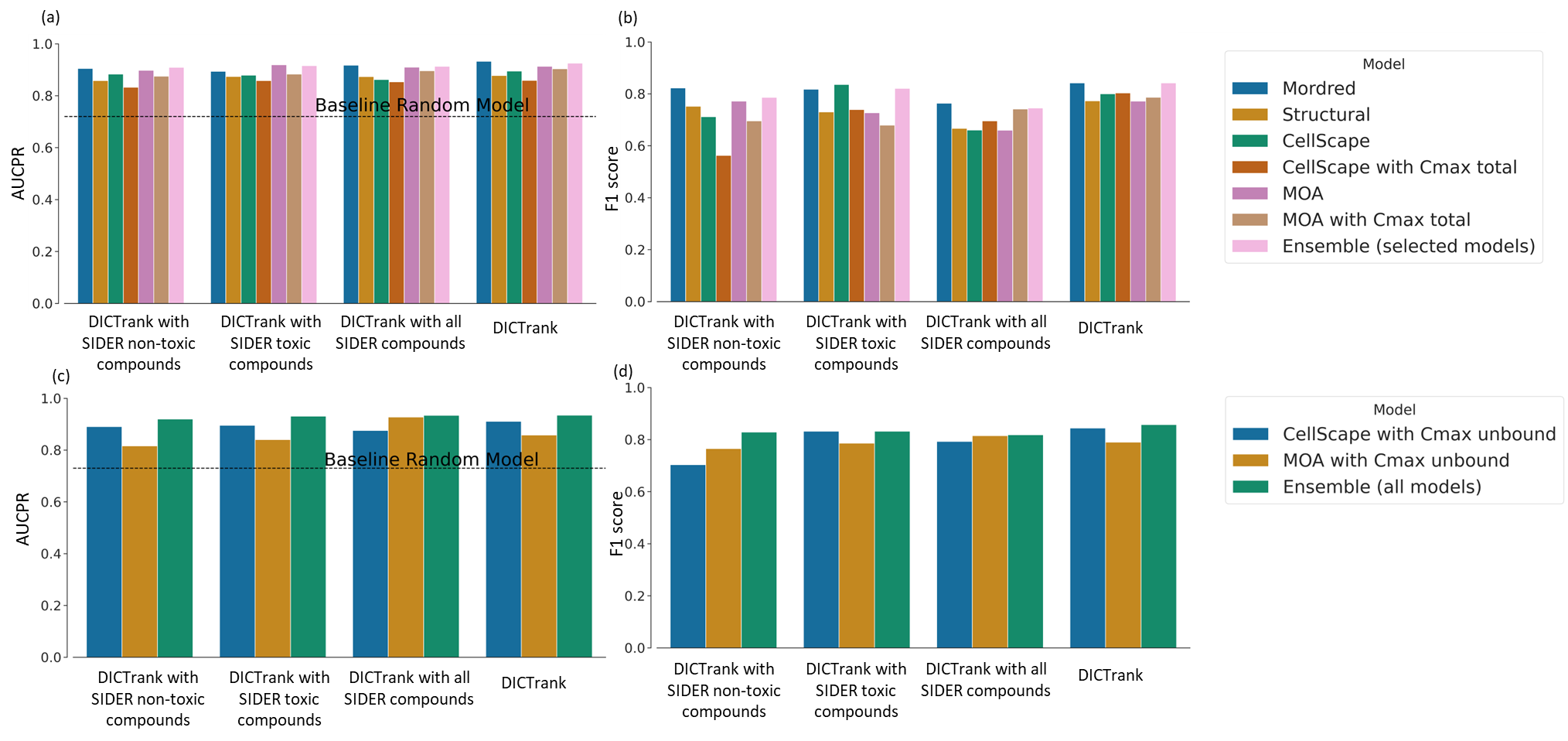

**Figure S3:** Evaluation metrics for models developed in this study using a larger external test set of 90 compounds evaluated using (a) AUCPR and (b) F1 scores, and for models using a subset of the external test set with 78 compounds evaluated using (c) AUCPR and (d) F1 scores
